# The Brain Knows enough to take into account Light and Shadow

**DOI:** 10.1101/2020.10.19.344838

**Authors:** Masahiro Yamamoto, Shinsuke Shimojo

## Abstract

Visual perception requires to infer object and light source color to maintain constancy. This study demonstrates the influences of environmental sunlight color trajectory (blue-white-yellow-red), and associated color of scattered light in shadows on color perception. In Adelson’s checkerboard shadow illusion, squares of equal luminance appear lighter or darker depending on whether they are inside or outside a cast shadow^1^. In some color variations, illusion magnitude is attenuated by specific colors of the cast shadow. Particularly in the green monotone environment (green checkerboard under green ambient and diffusion light), illusion magnitude reduces down nearly to zero. In contrast, shading by structure is not affected by the color environment. Thus, the cast shadow and shading by structure have distinct effects on surface color constancy. This illusion attenuation may be related to the absence of green in the natural environmental light spectrum, including in cast shadows. The brain may utilize the implicit learned trajectory of natural light to resolve ambiguity in surface reflectance. Our results provide a new formula not only to understand, but also to generate new variations of other illusions such as #The Dress.

---

The perceived color of an object is determined by two factors—its inherent light-reflecting properties and the color of the incident light. Thus, color perception requires the estimation of both object color and light source color^1,2,3^. For example, estimating the color of the light source is critical when the entire object or a part of it is in shadow. In a natural environment, the shadow color is dark blue due to the color of the sky and the Rayleigh scattering effect on longer wavelengths (long wavelength shielding effect)^2^. Thus, in shadows, an object is illuminated by a dark blue light source. To maintain perceptual consistency, the shadow color should be corrected according to the light source, even if not explicitly specified. In this study, shadows are treated as implicit ambient light, and the effects of lighting color on perception are examined. The study demonstrates that color perception involves adjustment based on the knowledge of the statistical wavelength distribution of natural objects and natural light, which may have been inculcated through experiences.

The perception of color and brightness depends on contextual cues and are influenced by cast shadows and three-dimensional (3-D) shape (shading by structure) (Fig. 1)^4,5^. Thus, the luminance image can be decomposed into three separate components: reflectance image, illuminance image, and cast shadow image (Fig. 1(b), (c), and (d)). Regardless of the context and 3-D shape, the pattern on the surface is perceived as stable. One cause of optical illusions is the brain’s adaptation to emphasize surface patterns, reflectance/texture, and 3D structure, regardless of variations in the lighting environment. Both shape images and shadow images contain contrast information, based on which color constancy is maintained. Color constancy under shadow arises from surface pattern recognition, which, in turn, requires eliminating the effect of light scattered in the shadow area. In other words, color constancy under shadow arises by identifying the surface reflectance/texture to remove the influence of the structure creating the shadow^6^.

**Figure 1.**
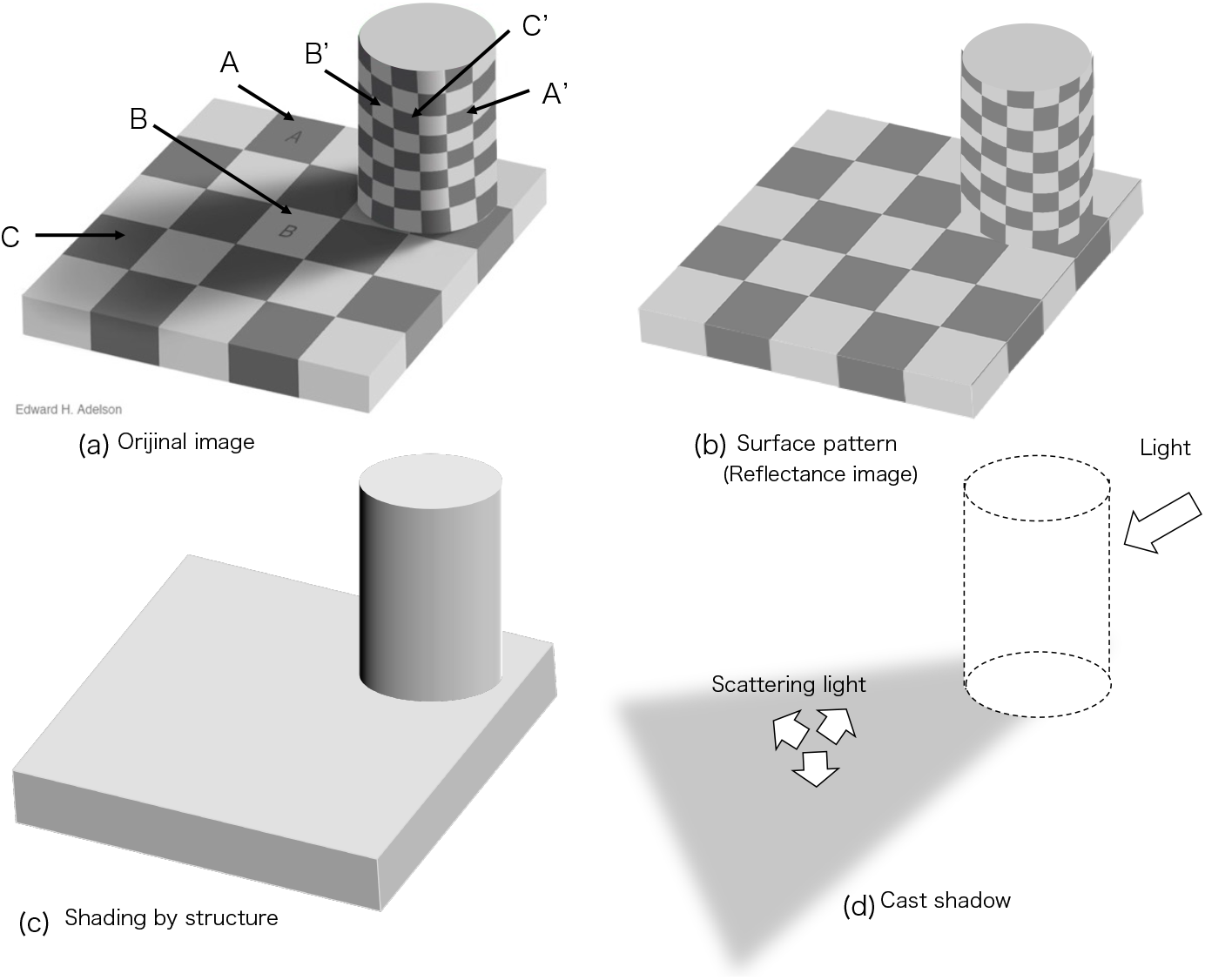
The “checker shadow” and its analysis by decomposition into three intrinsic images.^**6, 16**^. (a) Adelson’s original checker shadow image with checker shade on a cylinder is shown. (b)The intrinsic reflectance image, removing the lighting effect from (a), is shown. (c) The intrinsic shade image as an illuminance image without cast shadow is shown. (d) The cast shadow image alone is shown as an intrinsic shadow image of (a). Brightness perception is dependent on contextual cues. It is also influenced by three-dimensional shape. Furthermore, the luminance image into the eyes can be considered as the product of three other images: the reflectance image, the shade image, and the cast shadow image. And checker patterns are introduced as a high contrast pattern, so some very interesting phenomena occurs that is known as Adelson’s Checker Shadow Illusion. Irrespective of contexts and three-dimensional shapes, the pattern on the surface is perceived consistently, owing to the brain’s constancy mechanism, as expressed as: Visual image = Surface pattern × Shading(structure) × Cast shadow. One cause of our perception, sometimes leading to an illusion, is the brain’s adaptation to selectively extract or paying attention to the surface pattern irrespective of the lighting environment. Patches A, A’, B, and B’ have the same physical brightness. However, patches A and B are perceived with different brightness. Similarly, patches A’ and B’ are perceived as having different brightness. Since the brightness of patch B or B’ is located in the cast shadow or shade, the brightness is compensated and thus perceived as different brightness. For reference, patches C and C’ are physically dark, because patch A or A’ is located in the cast shadow or shade, and are actually perceived as dark.

Adelson’s Checker-Shadow Illusion is a famous example where the brain attempts to maintain brightness constancy under shadow based on the prediction of spatial regularity of surface texture for instance. In Adelson’s Checker-Shadow configurations, the observers typically perceive the luminance (color) of a part of the grid in cast shadow as brighter than the true physical brightness. In the most basic form, a gray square placed in an expected white square position, within a white–gray checker pattern, is perceived as white when under the cast shadow of a cylindrical object (Fig. 1(a) square B), even when the physical luminance is equivalent to the black-perceived square A outside of the shadow. The Checker-Shadow Illusion is created by recognizing surface texture (or reflectance) patterns and eliminating the influence of light scattered into the shadow region; on the other hand, the shade illusion is created by identifying the surface texture pattern and removing the influence of the structure. Further, the illusion can alter perceived hue (e.g., gray as red)^7^.

Usually, the shadow color is perceived as black, but the real color of the cast shadow is dark blue. If our color perception system knows basic light wave theory, i.e., scattering, then our implicit color perception system knows the real cast shadow color. Thus, the colored cast shadow caused an illusion in the form of system error, which we expected.

A computer vision uses complex calculations to handle cast shadows, whereas the human brain rarely mistakes cast shadows for surface textures. This is because the perception of a cast shadow is concerned with constancy and knowing its functions is a very important viewpoint in understanding constancy. We examined how perception changes when cast shadows are colored, which is possible in a very limited physical environment. Cast shadow perception is usually very robust, but does it remain so in artificial environments? It can easily be estimated that robustness leads to constancy.

We simulated a colored shadow environment and filled the space with colored ambient light. Then, the checkerboard was colored the same as the ambient light color. The cylinder was placed as a shield on the board and color illumination that consists of ambient light color and white color was illuminated from one direction of the cylinder. The result is a colored Adelson’s checkerboard with colored cast shadows over the same colored checkerboard. We found that in some color versions, the illusion of the brighter checker was extremely attenuated by specific colors of the cast shadow. A typical observer experiences vigorous illusions as shown in Fig. 2; however, a much weaker effect of some color versions were recorded at 12, 168, 183, and 202 degrees on L-M vs S-(L+M) plane in Derrington, Krauskopf, and Lennie (DKL) color space as illustrated in Fig. 2^8,9,10^. Indeed, when the ambient color, diffusion light color, and checker color are green or purple, the magnitude of the illusion, despite being lighter B than A, either intuitively or quantitatively (as later) is nearly zero. The first section of this study explains this optical illusion as a correction effect for the constancy of the cast shadow color. Shadows can be considered ambient light when no explicit light source is specified. Therefore, the first experiment in this study also examines the color constancy in the absence of explicit light source information.

**Figure 2.**
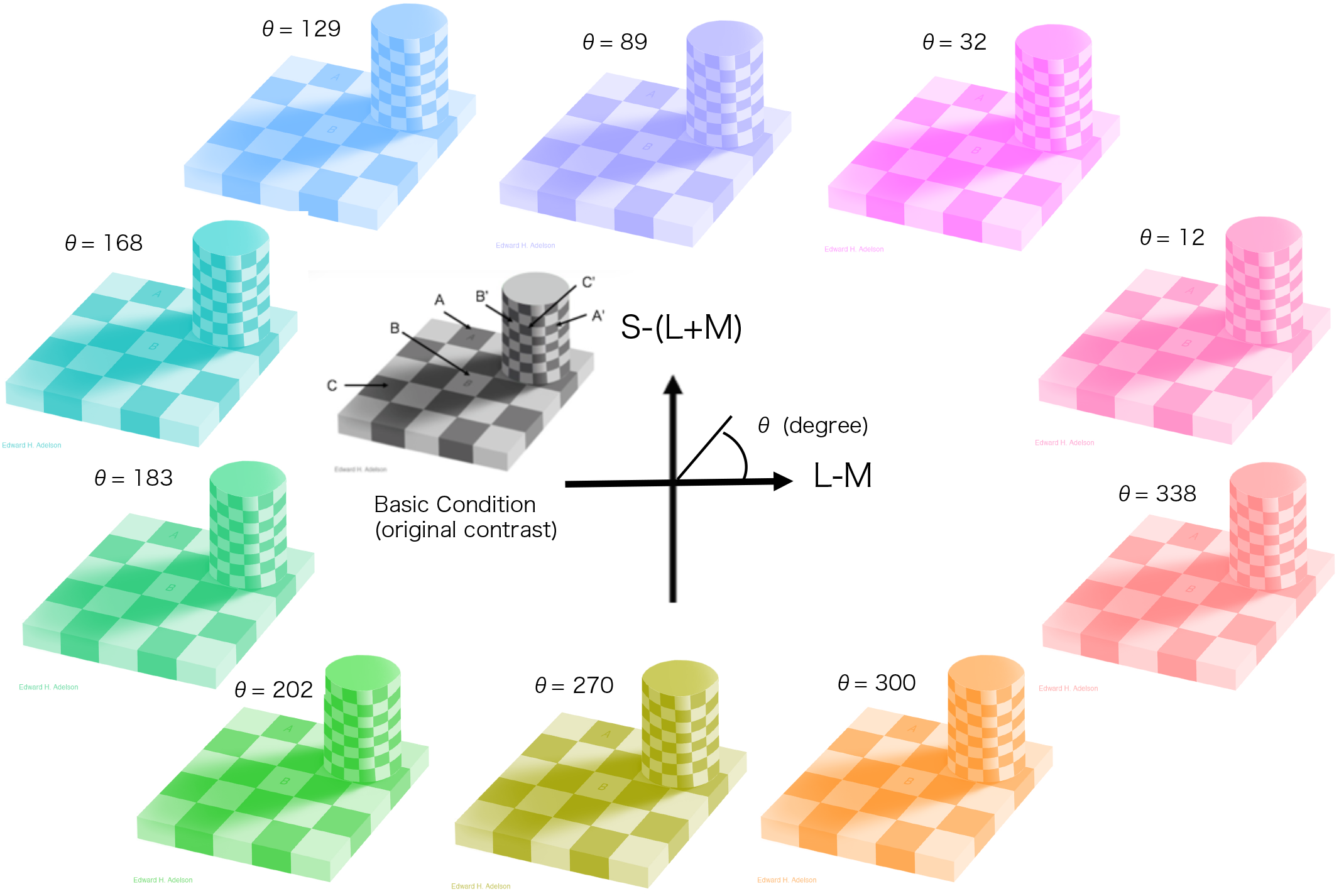
Example of stimulus display used in the contrast ratio 0.13 condition. This map shows the angle of stimuli in the Derrington, Krauskopf, and Lennie (DKL) color space.(24,25,26) The DKL color space in cone space is constructed using the opposite color in the lateral geniculate nucleus. Its hue axes represent constant L-M values and content S-(L+M) values (L: long, M: medium, and S: short-wavelength cone sensitivity). Experiment 1, in which square B on the board is changed to be equal to square A on the board, is a measure of the illusion magnitude in the cast shadow. Conversely, in Experiment 2, in which square B’ on the cylinder surface is changed to be equal to square A’ on the cylinder, is a measure of the illusion magnitude of shade condition. The squares A, B, A’, and B’ are shown in the basic condition diagram (the black-white image). The same notation applies to images of each color condition. However, in the shade experiment (Experiment 2), cylinder parts were spatially enlarged to align the area of the square with the cast shadow experiment. (See Supplement 1 for basic generation rules) If the image is enlarged and observed as a large stimulus, the illusion can be easily confirmed.

Such color specificity in the shadow illusion raises the question—whether this specificity is also observed in color versions of other brightness illusions. We hypothesized that the effect of the shadow illusion is strongly influenced by the shadow context; thus, we examined the influence of shadows containing contrast information in the second part of this study. To this end, we examined a checker shade illusion constructed by wrapping the cylinder of the Adelson’s Checker Shadow image with a checker pattern of several colored ambient light patterns. Therefore, there are two illusion groups in this paper: the cast shadow illusions group and the self-shading illusions group based on a 3D structure. The original Adelson’s Checker Shadow Illusion belongs to the cast shadow illusion group. Our first proposal is that the color version of the cast shadow illusion was attenuated green.

We also propose that the color perception is influenced by visual adaptation to the color distribution of natural scenes and investigate how natural color statistics change under sunlight. It is conceivable that color perception arises during evolution or development by implicit learning of the natural environmental color distribution, particularly the natural daily trajectory of ambient light. Therefore, in the third experiment of this study, we applied light source estimation to the “#The Dress” image, which is perceived as white and gold or blue and black due to the estimation of two unclear light sources^11^. We propose a new interpretation of “#The Dress” in the context of the change in the physical color distribution of natural objects caused by daylight trajectory, which is consistent with our interpretations of the Adelson’s and other illusions, and, clarifies the mechanism based on natural scene color distribution simultaneously. Indeed, we construct another color variation of “#The Dress” for verification.

Two experiments provide the bases for these conclusions; sunlight is the first choice for lighting color and object colors are basically based on the natural color statistical distribution, where the functions are independent of shape of the object. First, the participants evaluated the cast shadow illusion magnitude by adjusting the statistics of a target (e.g., in cast shadow) to match a comparator (e.g., outside a cast shadow). Second, the participants evaluated the illusion magnitude of the colored shade illusions. We, then, conducted two experiments using Adelson’s type illusions, comparing shadow/shade illusion magnitudes from color combinations within or outside the color statistics of natural scenes. Finally, we demonstrated new forms of “#The Dress” based on the proposed influence of natural light statistics on the color perception.

## Method

### General Method

The purpose of the present series of experiments was to determine whether color variations of shadow and shade influence illusion magnitude by CIEDE2000^12^. Experiment 1 demonstrates that color variation influences Adelson’s checker shadow illusion, and Experiment 2 investigated whether color influences the shade illusions (shade by structure). Furthermore, to better understand human color perception, this study qualitatively examined the differential effects of natural color scene statistics with sunlight and in cast shadows.

### Measurement of the Illusion Magnitude

Before conducting the tests, the protocol was approved by the institutional review board and written informed consent was obtained from all participants.

The two experiments studied 12 females and 14 males aged 18–45 years, 16 (eight females and eight males; age range: 19–35 years, average = 23.8 years; sd = 5.3) in Experiment 1 and 10 (four females and six males; age range: 24–36 years, average = 30.2 years; sd = 3.9) in Experiment 2. They were naive about the experiment, and all had normal or corrected-to-normal vision.

### Stimuli

Figure 2 shows the basic stimuli used in Experiments 1 and 2, consisting of a checker pattern and cylinder. Details of the color conversion rules and physical interpretation are described in Supplement 1. The color image variants (Figure 2) were created by changing only one RGB component to the maximum value (255). Images created by this method all contain virtual colored shadows (see Supplement 1 for details of the color conversion rule). We calculated the luminance of the brightest and darkest squares in the DKL color space and determined the contrast ratios^34^. Subsequently, we changed hues while maintaining constant luminance (lum) contrast ratio, where

Contrast ratio (Cr) = (lum-brightest − lum-darkest) / (lum-brightest) + lum-darkest). All versions were constructed for assessing the luminance of the brightest area. Hues were calculated as the central angle of the <L − M> vs. <S − (L + M)> plane in the DKL color space.

The method of adjustment was used to measure illusion magnitude in each experiment. The participant adjusted the color luminance of square B in the cast shadow to be equal to that of square A in the brighter region using a computer interface slider (see Supplement S2 for details). The illusion magnitude was defined as the difference between the adjusted color luminance of square B and the original color luminance of square B. Stimuli were presented on a color monitor in a dark room. A maximum illusion magnitude (=100) was obtained when the color luminance of square B after adjustment was equal to the color luminance of the brightest area in the checkerboard (square A). The computer screen subtended 56° horizontally and 38° vertically. Three contrast conditions were tested: (i) Cr = 0.13, (ii) Cr = 0.27, and (iii) Cr = 0.36. The entire checkerboard extended 42° × 28° on the monitor.

### Physical Measurement of Color Distribution

Visual input consists of the light source and reflections from objects, so the final spectral distribution projecting on the retina is critical in color perception. However, color perception is also influenced by the natural color statistical distribution^13^. We examined shadows as well as natural color distributions to provide support for our conclusions about shadow perception.

The color distributions of natural scenes illuminated by reflected light only were measured using a small-sized spectrometer (Hamamatsu Photonics, C12666). Each measurement was corrected against a white standard sheet. Following previous studies, selected images were of natural objects such as flowers, leaves, and trees reconstructed in DKL color space along three post-receptor axes, L + M, L − M, and S − (L + M).(24-26) Furthermore, the distribution was calculated by comparing it with the reference data^14^.

### Experiment 1 Colored checker shadow illusion Design

The participants were presented the colored stimuli (Figure 2) in random order. Moving the slider left or right changed the color luminance of the target square (0–255). The task was to adjust the color luminance of square B so that it was perceived to have the same luminance as square A. Results were logged for analysis by double clicking on the final adjusted image. The color of the rectangle was programmed in JavaScript. The adjustment color luminance was changed smoothly between the brightest and the darkest (see Supplement 2). Initially, the slider was randomly placed. The illusion intensity was subsequently calculated from the contrast ratio between the target square and the surrounding dark lattice. Gamma-corrected stimuli with resolutions of 1240 × 1024 pixels and color resolutions in sight bits per channel were presented on an Eizo L767 monitor driven by an Intel HD Graphics driver installed on a Toshiba Dynabook Satellite K45 266E/HD. The monitor was placed 40 cm from the participant.

### Experiment 2 Colored checker shade illusion Design

The observer was presented with randomly colored stimuli and was requested to adjust the color luminance of square B by moving the slider below the figure until it appeared to be the same color luminance as that of the reference square A. The initial position of the slider was randomly set. Measurements were performed twice for each image. The primary method was the same as that followed in Experiment 1. The difference is that in Experiment 2, the subject manipulated the target square on the side of the cylinder in the shade. In Experiment 2, cylinder parts were enlarged to align the area of the square with that in the cast shadow experiment (See Supplement 3).

### Statistical Analyses

Differences of illusion magnitude between stimuli were analyzed by two-way or one-way analysis of variance (ANOVA). All statistical analyses were performed using EZR (Saitama Medical Center, Jichi Medical University, Saitama, Japan), a graphical user interface for R (The R Foundation for Statistical Computing, Vienna, Austria). More precisely, it is a modified version of an R commander designed to add statistical functions frequently used in biostatistics.

## Results

### Experiment 1 Colored cast shadow illusion

Fig. 3(a) presents the relationship between illusion magnitude and color for Adelson’s checkerboard shadow illusion (detailed results are shown in Supplements S4). To eliminate the influence of contrast, the experiment was conducted with three contrast conditions. Two-way ANOVA indicated significant main effects of both color (F(9, 17612) = 9.9730, P < 0.001) and contrast ratio (F(2, 4389) = 11.1836, P < 0.001) but no significant interaction (F(18, 4363) = 1.2353, P = 0.2281). Changing the hue while maintaining the contrast ratio altered the illusion magnitude (filled circles), with a significant magnitude decrease in the green shadow environment (Green Region, Fig. 3(a)). At the lowest contrast ratio of 0.13 (shown in Fig. 3(a)), the green illusion intensity became CIEDE = 2.82, which is below the general color difference discrimination limit of CIEDE = 3.5, effectively eliminating any illusion^15^. One-way ANOVA for each contrast condition revealed that the DKL color effect was significant at Cr = 0.13 (F(6,105) = 9.338, P < 0.001) and Cr = 0.26 (F(6,105) = 2.308, P = 0.0392) but not at Cr = 0.37 (F(6,105) = 1.336, P = 0.248). We also performed Bonferroni correction to ascertain which colors were more effective and found that blue and green areas differed significantly in illusion magnitude (see details results in Supplement 4).

**Figure 3.**
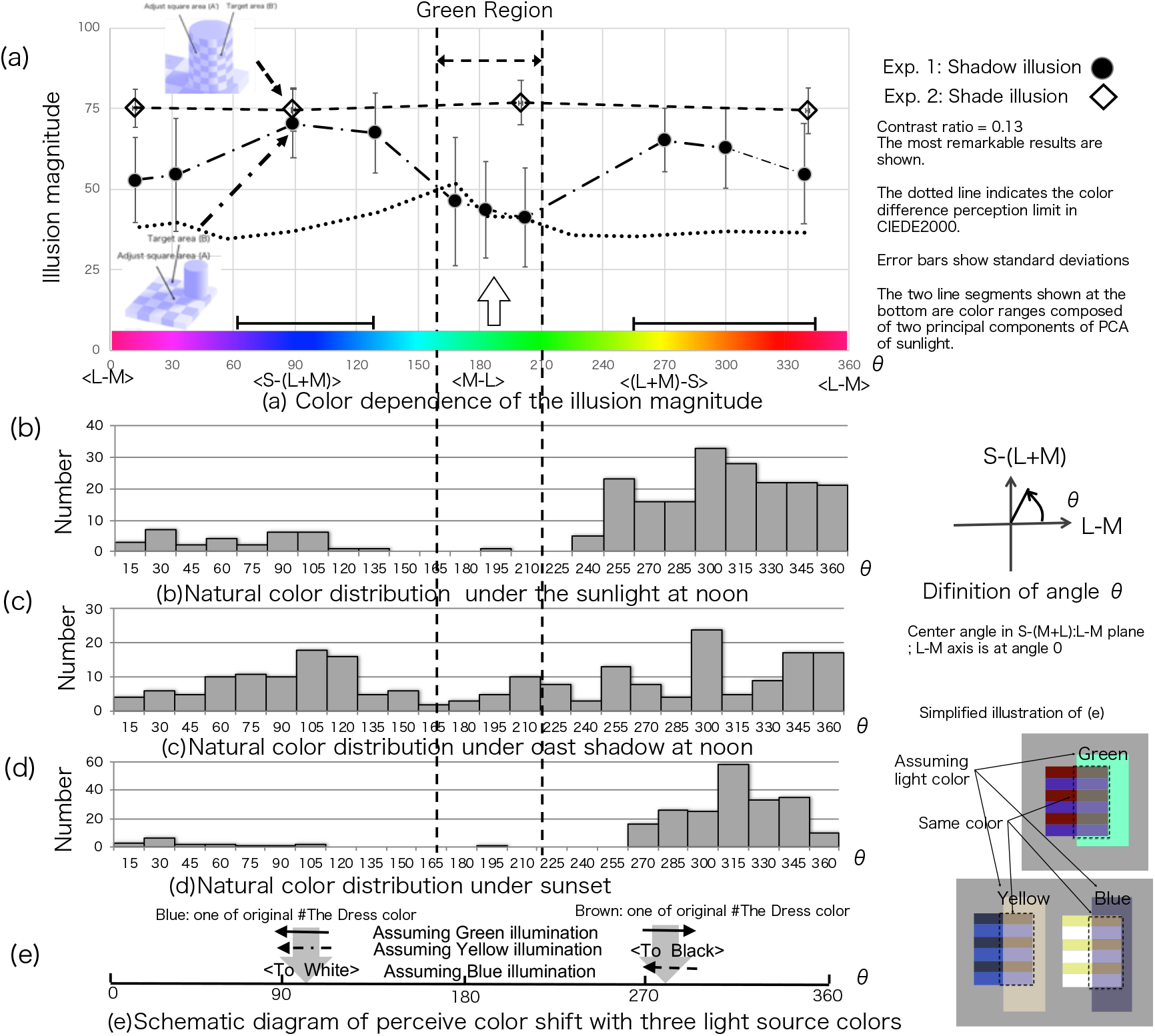
All results derived are summarized; cast shadow illusion and shade illusion, and the distribution of natural scene color. Fig. 3(a), which presents two graphs of illusion magnitude against color angle, circles show the results on shadow illusion, and diamonds show the results on shade illusion. In Experiments 1 and 2, the color of the target square was adjusted to the same color as the reference, and the illusion magnitude was measured, see the method section for more details. The illusion magnitude 0 is the case in which no illusion occurs. The illusion magnitude 100 is the case in which the color of the target is the same as that of the darkest square in the figure. The color difference was calculated using the ΔE00: CIEDE2000 color-difference formula^12^. The dashed curve shows the different color perception limits in CIEDE2000^15^. This factor is the low limit of color difference for naive observers. In the region of the green [DKL angle 129,168, 183 degrees: denoted by a white arrow in Fig. 3(a)], the illusion magnitude is either lower than or equal to the color difference limit. Posthoc analyses with a one-way ANOVA on condition of contrast ratio = 0.13 that show the attenuation of the illusion magnitude of checker shadow depends on the DKL color angle. In the green region, color difference conditions were statistically significant. A one-way ANOVA for color indicated a significant effect F (9, 15197) = 7.993, P < 0.001. Natural scene statistical data are shown below: (b), (c), and (d). The color distribution was created by measuring and adding data calculated based on the reflectance dataset^14^. (b) Natural color distribution in the sunlight at noon is shown: (c) Natural color distribution in the cast shadow at noon is shown: (d) Natural color distribution at sunset is shown. Attenuation of the shadow illusion depends on the natural color distribution in the sunlight, and cast shadow also depends on principal component analysis (PCA) of the natural light source made by the sunlight and the blue shadow. Two main components of PCA results of the sunlight spectrum including cast shadows are shown at the bottom of Fig. 3(a)^50^. The two main components can explain the changes in (b), (c), and (d). (e): Illustration of the perception rule used by our calculation of Kullback–Leibler (KL) divergence^36^. The vertical gray arrows denote stimuli used in the original #The Dress illusion. Horizontal arrows indicate a perceived color shift with three different illumination colors. If either blue or yellow illumination is assumed, one side moves to the coordinate origin, and it is not shown. Right-hand side small color images are these rather interpretations for variations of #The Dress illusion, assuming luminance color as green, yellow, and blue. The results of the measurement of illusion magnitude are shown in detail in Supplements 4 and 5. The details of the calculation are shown in Supplement 6.

### Experiment 2 Colored shade illusion

Fig. 3(a) also illustrates the effect of color on the shade illusion intensity at Cr = 0.13 (see details in Supplement 2). Neither contrast nor color significantly altered shade illusion intensity (open diamonds). Two-way ANOVA indicated no significant main effect of color (F(3, 94) = 0.8281, P = 0.4812) or Cr (F(2,175) = 2.30684, P = 0.1045) and no significant interaction (F(4,220) = 1.3662, P = 0.2530) (see Supplement 5 for details).

### Natural scene color distribution

We speculated that this effect of color on illusion magnitude may stem from brain compensation (or adjustment) and that this adjustment is learned implicitly based on the natural spectral distribution of sunlight, objects, and shadows in the natural environment. The color distribution of natural objects in the DKL color space has been examined in previous studies of visual chromatic adaptation. The DKL color space consists of three post-receptor axes [L + M, L − M, S − (L + M)], which are the basis for color vision. Natural color scene statistics were used to reconstruct the contrast along these three axes in the DKL color space. We also selected the DKL color space and examined the effects of cast shadow colors.

The color distribution of natural objects was calculated based on 200 reflection spectra^14^ by measuring sunlight and shadows during the day. For 40 data samples, the colors of natural objects were measured and adjusted (corrected). Fig. 3(b), 3(c), and 3(d) show the color of a natural scene object plotted in shade, in shadow at noon, and at sunset, respectively.

Object colors have a biased color distribution along the daily sunlight trajectory in the DKL color space because sunlight changes from white to yellow to red as the day progresses. The coordinate range of DKL color space is 90°, and the range is from 270° to 330°. The coordinate range of ambient light (scattered/diffused light) in the cast shadow is also about 90°. The color range calculated by principal component analysis from the sunlight trajectory^16^ (including shadows) is displayed as a line below Fig. 3(a).

Changes in the histogram over the calculated color range are observed for Fig. 3(b), (c), and (d). That is, Fig. 3(b), 3(c), and 3(d) indicates how the natural color distribution changes with natural light. In Fig. 3(b), (c), and (d), the green region crossing the graph up and down have no light source and very little object color distribution.

### Application of findings to #The Dress

Fig. 3(e) shows our interpretation of the well-known ‘#The Dress’ image. The original coordinates of #The Dress, dark blue and dark brown, are 91° and 274° on the S − (M + L) vs. L − M plane in DKL color space. Of the two peaks in the natural color distribution during the daylight, one is less than 91° and the other is larger than 274°. If the original is assumed, green lighting can superimpose the two original peaks of #The Dress color onto the same peaks of the natural color distribution. In other words, stochastically, the original two coordinates may include green illumination. However, our cast shadow illusion experiments show that green environment color selection probability is almost zero when no illumination is explicitly shown. Therefore, only one of the two coordinates is selected, with one of two luminance colors calculated in #The Dress.

## Discussion

The visual system uses various local and global rules to reconstruct scenes and maintains constancy under changing light conditions^17-22^. For instance, the difference in brightness and edge ambiguity actively contribute to shadow recognition. Colored shadows (except dark blue and red) are extremely rare in natural environments, and, therefore, the visual rules related to shadow color recognition can be revealed by colored shadow illusions. Two predictions can be made if our interpretation of Adelson’s green attenuation is correct. First, the green color attenuation should not be observed in shade illusion (as opposed to cast-shadow illusions). Second, based on the natural statistics and the brain compensation rules that we discovered, we should be able to generate new illusions, such as variations of “#The Dress.”

In Experiment 1, we found that the magnitude of the illusion created by colored shadows, in variation of Adelson’s checkerboard shadow illusion, is highly dependent on the particular color of ambient light (Fig. 3(a), closed circles), with almost complete attenuation using green ambient light. The differences in brightness and ambiguity of edges are also observed in shade conferred by the 3-D structure, which may not be the cause of the attenuation. Therefore, unlike the shadow illusion, there was no color dependence in the shade illusion magnitude, as shown in Experiment 2 (Fig. 3(a), open diamonds). The detailed descriptions of the illusory effect of shade have been reported in previous studies^23-28^. We confirmed these results and further showed that there is no change in the illusion magnitude across the color variations related to a cast shadow. By structure, cast shadow and shade have different effects on the constancy of surface colors. This difference also suggests that perceptual mechanisms can distinguish between a shaped shade and a cast shadow even when stimuli are similar.

### Influence of Natural Scene Color Statistics

The light source in a natural environment (usually sunlight) should have a strong influence on our perception of shadows (Fig. 3(b), 3(c), and 3(d)). Indeed, our spectrometry measures revealed that the colors of objects in a shadow show a sharp blue shift as compared to the color under sunlight. We also found that the blue-green light is low in intensity throughout the daily sunlight trajectory and thus contributes little to the spectral distribution of natural objects and shadow color. Thus, the color specificity of the illusion attenuation may reflect brain compensation (or tuning) to the daylight spectral distribution in a natural environment.

The color distribution of natural objects in DKL color space was also described in a previous research on visual color adaptation to the color distributions characteristic of natural images^13^. For instance, the overlap of the sunlight trajectory and the color distribution of natural objects has implicated the adaptation of color perception. Previous reports show that the color distribution of natural objects is related to the trajectory of sunlight in color spaces^30-32^. The color exits a distribution of natural objects in the DKL color space, which is formed by related sunlight. As a result, the distribution is shaped to be very low in the green and purple regions, which are not in the sunlight trajectory^13,29^. Our idea is that the shape of the color distribution can be a normalization of the color distribution of natural objects concerning sunlight; we believe that this normalization makes the cone difference more efficient during color perception^33^. In other words, sunlight is specially tuned as a light source, which means that green light sources, which do not occur as a trail of sunlight, have strange properties. In the absence of any explicit illumination color information, the only way to perceive an object color is to use natural illumination, sunlight, and the color distribution of natural objects.

Using the color distribution of natural objects to determine an object’ colors means calculating the overlap between the color distribution of the object and the distribution of the natural objects. In case of a roughly white-colored object, we do not misread white in sunlight from white to red, even if it is in the cast shadow. In other words, the color distribution of natural objects is considered to reflect the estimated probability of the light source color for the achromatic color of objects. This means that green, which has almost a few distributions, is unlikely to be a light source color. Indeed, the attenuation of cast shadow illusion occurs within the band (green light) with an axis perpendicular to the trajectory of sunlight, the distribution of natural objects, and the color distribution of shadows (Fig. S6-1). Under natural conditions, green light color is very rare, so the attenuation of the illusion occurs primarily at this low intensity.

### The Context of Cast Shadow And Shade

Unlike the cast shadow illusions, the shade illusion occurs regardless of the source of light. Different illusions can be obtained by simply rotating a stimulus on the image in a green environment. In this case, only the changing shade and shadow, due to rotation, promote the illusion. A new illusion involving the switching between cast shadow perception and shade perception is shown in Fig. 4, to confirm this context-dependence. A typical observer experiences vigorous illusions (Fig. 4(a)-I, II, 4(b)-I, II, 4(c)-I, and 4(d)I, II) but with a much weaker effect (Fig. 4(c)-II). An automatic perception system consisting of light from nature is used for the illusion. The rectangle in the image is connected to the target grid, and therefore, a gradation appearing in the rectangle is an illusion. When the cylinder is turned sideways, with the assumption that the light shades from the top, the checker pattern under the cylinder will be recognized in the shadow of the cylinder. As a result, the illusion due to the vertical shadow disappears. In Fig. 4, the difference between cast shadows and shades occurs only in the green environment. The example of optical illusion strongly supports our hypothesis that the context of light color is automatically estimated from the color trajectory of sunlight without the information of source light color explicitly (other variations of the optical illusion are shown in Fig. S7).

**Figure 4.**
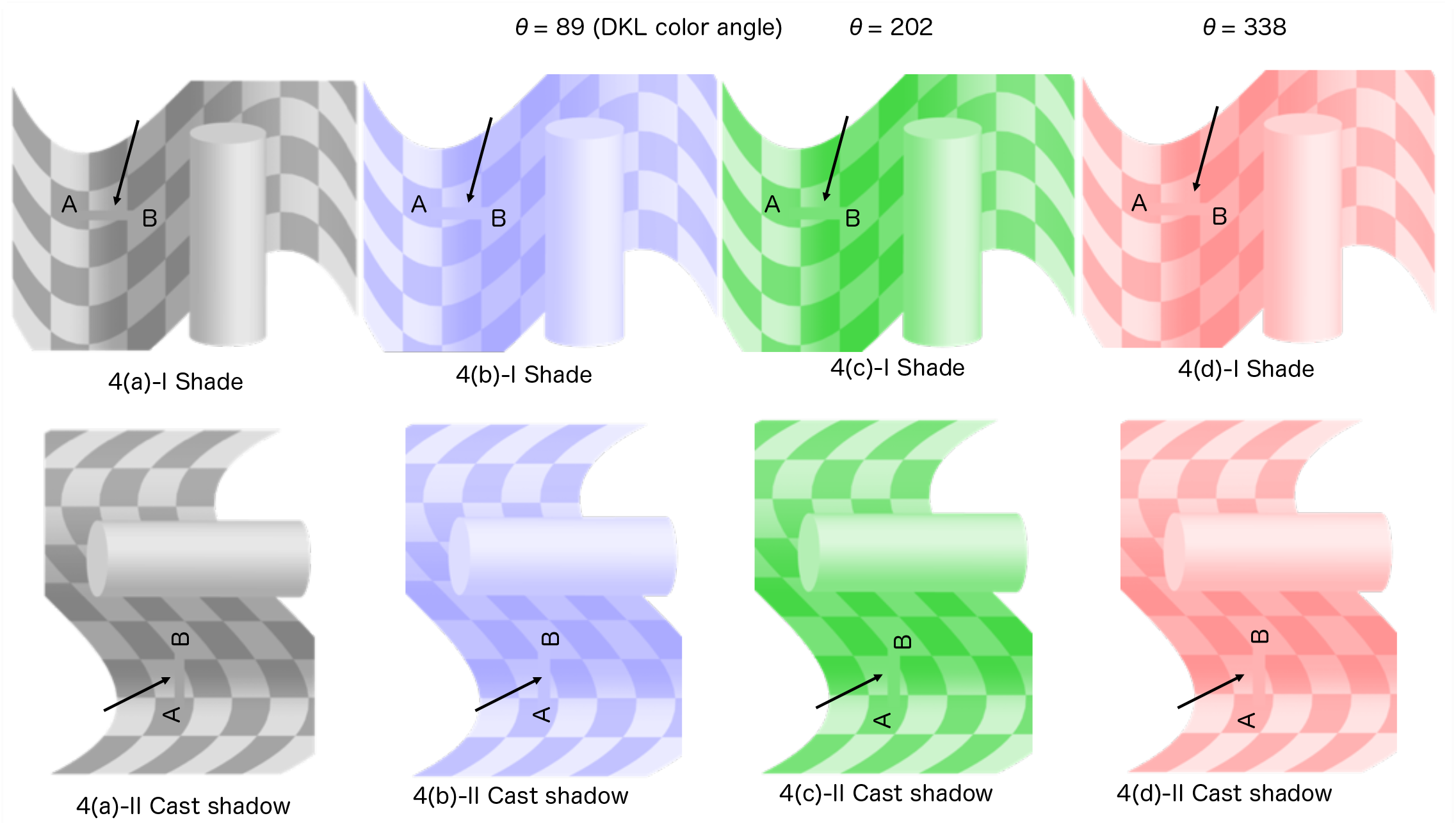
Illusions created by the subjective perception of cast shadows and shades under several ambient colors. Assuming our theory is correct, the cast shadow illusion is different from the shade illusion at the green. We created two meanings by rotating the picture. Images with the suffix ‘I’ have a vertical cylinder. Though squares A and B are physically the same, B seems to be much brighter than A. Square B is in the shade with the snake shape of a checkerboard. Images with the suffix ‘II’ have a horizontal cylinder. In every suffix II image, B also seems to be much brighter than A in general except for 4(c)-II. Square B is in cast shadow of the horizontal cylinder, helped by the fact that it is lit naturally from above. In both I and II, the illusion makes the gradation on the rectangle bridge connecting A and B as the arrow position. However, this gradation is only different in Fig. 4(c)-II. In Fig. 4(c), the illusion effects result from the difference in perception of shadows (II) and shades (I). The illusion magnitude is attenuated at Fig. 4(c)-II. We believe that most observers would see A and B in a more similar colors (brightness) than in all Fig. 4-I color variations (including the green Fig. 4(c)-I), and all the other 4-II color variations (a, b, and d). Another type of shadow/shade illusion is shown in Supplement 7, Figure S7.

### #The Dress

In general, if the illumination color is known, color constancy is achieved simply by shifting the illumination color. As a result, the object color should become close to the inherent probability distribution. We combined this principle with illuminant color estimation when not specified. Thus, if the illumination color is unknown, color constancy can only be established by statistics of color (both the illumination color and object color) based on our experiences. In other words, the brain should select a common light source (usually daylight) for illumination, estimate colors of objects within that light source, and compare it to its inherent color map. The object color is then determined by maximizing the probability of the source illuminant color and the object color estimation. Our cast shadow illusion results suggest that the light source color should be along the trajectory of sunlight or the color of the shadow from sunlight should be considered for estimation of the light source.

The above hypothesis may explain the large individual differences (mostly two types among people) on color perception in “#The Dress,” where the lighting information is not explicitly given, and thus, inferred together with the reflectance (or the appearance of the material surface) from various surrounding cues. Therefore, we examined whether our theoretical framework also holds in such a situation and attempted to predict or produce a new variation of “#The Dress.” This variation is meant to be an additional test of our interpretation, i.e., when the information on lighting is not explicitly given, the light source color estimation should be along the sunlight trajectory or shadow color.

Several reports state that “#The Dress” is one of the hallmarks of internal computation in the visual system; a principled view to quantify the internal computation has not been proposed yet. It explains the probabilities of color selecting activity as a measure of the distance of sensory input color data and statistical color data on the specified color feature space using Kullback–Leibler (KL) divergence^33,34^. The KL divergence calculation is often used widely as a good measure of the distance between two probability distributions. And given that neural firing follows Poisson distribution, it is natural to use KL divergence as a standard statistical method with logarithms to compare them^35-38^.

And if our hypothesis is correct, we can create another color version of “#The Dress” illusion. Fig. 5(a) shows the color combination of the original “#The Dress” illusion, which has emerged from blue and black under light yellow illumination (left in Fig. (a)), or white and yellow under cast shadow (we would not call it gold, the usual color perceived because it will be golden if the yellow region has a specific contrast distribution^39^). The rules that have stemmed out of our hypothesis allow us to determine the best color combinations to perceive an object color that minimizes the distance of an object color distribution and the natural color statistics in the DKL color space while eliminating green or purple as an illumination color. The object color estimation and illumination color estimation were both probabilistic problems based on our experience of making the database of perception.

**Figure 5.**
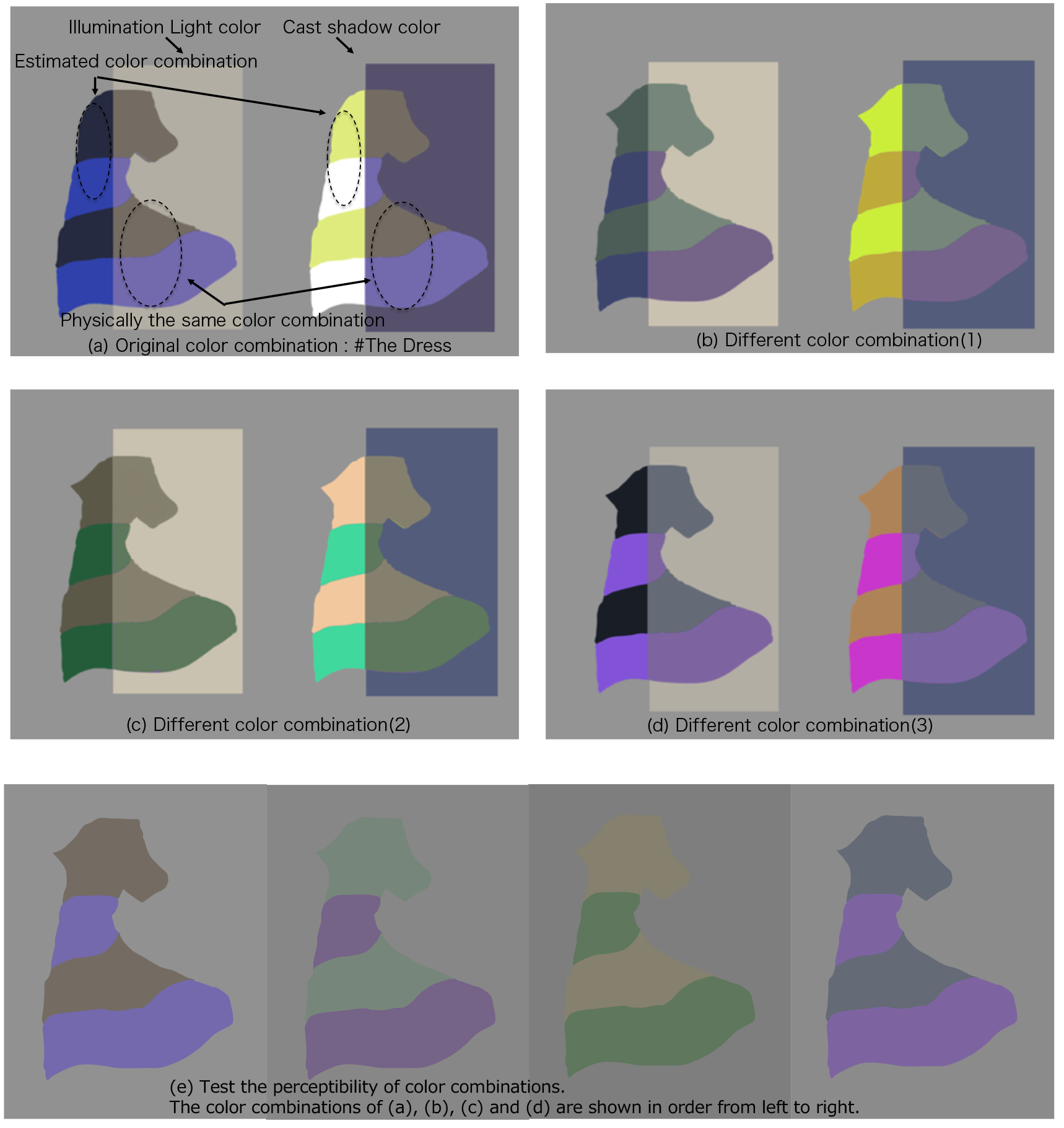
Different color variations of “The Dress” type illusion based on our hypothesis and generation rules. Our cast shadow illusion results suggest that the light source color should be along the trajectory of sunlight or the color of the shadow from sunlight should be considered for estimation of the light source. Therefore, we examined whether our theoretical framework also holds in such a situation and attempted to predict or produce a new variation of “#The Dress.” All combinations are calculated using the same rule, which is to maximize the probability of existence in the DKL color space while not choosing green and purple as an illumination color. The KL divergence is calculated correctly as a probability of combination color in the DKL color space, and the certainty of the estimated illumination color is also calculated by the KL divergence. Those are equivalent to the Wasserstein metric with the estimated combination colors, where the illumination color is a constraining factor. Two-color of the object, e.g., the dress color of #The dress, is plotted in the DKL color space. Then, we calculated the KL divergence to move two plotted color points as maximum overlap natural color statistical data. We found that the moving direction for the minimum KL divergence is either green or purple direction, which is not acceptable for natural light. Finally, the combination color was found with only a sunlight trajectory color (See Supplement 6 for an overview of the calculations). (a) The color combination of the original “#The Dress” illusion is shown. The perceived color combination has arisen from a color combination of either blue and black under cream-yellow illumination [left in Fig. 5(a)] or yellow and white under a cast shadow (right in Fig. 5(a)). [We would not call it “gold” here, as it will be golden if the yellow region has any specific contrast distributions^38^]. We show (b), (c), and (d) as other color combinations. (b) The perceived color combination has arisen from a color combination of either dark green and dark purple under cream-yellow illumination (left in Fig. 5(b)) or yellow and ocher under cast shadow (right). (c) The different color light sources reveal a color combination of either dark gray and dark green under cream-yellow illumination [left in Fig. 5(c)] or apricot and cobalt green under a cast shadow (right). (d) We can also perceive two different color combinations with the same physical color combination only, i.e., a color combination of either black and purple under cream-yellow illumination [left in Fig. 5(d)] or tan and ruby red under cast shadow (right). Fig. 5(e) shows the combination of the selected two colors when the background is achromatic. The background color is matched to the gray in the middle of the two colors’ brightness. It may be difficult, but you may be able to experience two interpretations of the color combinations in each of (a), (b), (c), and (d).

Therefore, in our hypothesis, there are two steps for the color calculation rule for basic color perception without explicit light source information. In the first step, we search the light color that is maximized by the probability that the target two colors overlap the natural color statistics distribution by the KL divergence. And if the light color is selected from a green (or purple) light source, the second step is needed because our color estimation system cannot choose the green light color without explicit light source color. In the second step, we search the light source colors with sunlight color only, from dark blue at cast shadow, white, yellow, to red, which maximizes the probability that the target colors overlap the natural color statistics. The two KL divergence are equivalent to the Wasserstein metric if we substitute estimating the combinatorial color using an illumination color as a constraint factor^40^.

The most probable combination of light and object colors is selected by the above two steps. In a special case, as “#The Dress,” two lighting colors are selected based on equal probability. Thus, two perceived color combinations can be generated by two other light source colors as an equal probability. There are new color combinations as type “#The Dress,” Fig. 5(b), (c), and (d), those were calculated with our two-step rule. Our visual system knows how shadow reflects light, so the object colors are determined with a simple estimation rule of light and shadow color.

In general, when determining colors in an environment where the illumination information is not explicitly given, it is necessary to maximize the probability of the perceived colors from the probability distribution of object colors and that of light source colors. The associated constraint was indeed observed as the color dependence of the case shadow illusion magnitude in our results (Exp.1).

There have been several theoretical interpretations offered on “#The Dress.” For example, Brainarad et al. (2015) analyzed the selection of the anchoring point of the light source and the overall color shift^41^. Although our results are based on statistical methods, they are consistent with previously reported interpretations^42-50^. Our interpretation also demonstrates that an objective, quantitative optimization (for the illusion) is possible based on the illuminant color estimation specificity and statistical background. Indeed, eye inspections of Fig. 5 indicate that our method was successful in producing new “#The Dress”-type illusion color variations.

## Conclusion

We proposed that the brain uses the probability distribution of natural light and object colors that are derived from experiences, to facilitate veridical perception. This study suggests that the human color vision system utilizes the features of sunlight, including cast shadow color and natural scene statistics for optimized perception. Indeed, the brain knows light and shadow to determine the color of objects and luminance.

## Acknowledgement

We thank our colleagues from TOSHIBA R&D center who provided insight and expertise that greatly assisted the research, although they may not agree with all of the interpretations/conclusions of this paper.

## Notes

### Competing Interest Statement

The authors have declared no competing interest.

