## Supplement for "The Brain Knows enough to take into account Light and Shadow"

### Supplement 1 Making rule and physical properties of stimuli.

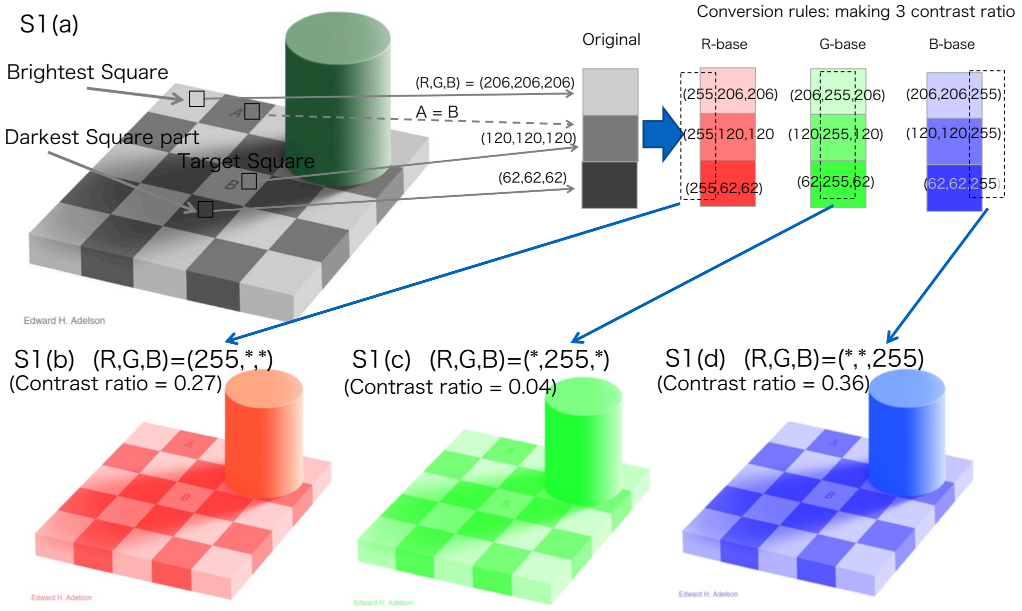


Figure S1. The color conversion rule of stimuli.

Figure S1(a) shows the original Adelson checker shadow illusion, and Figs. S1(b), (c), and (d) are variations with colored shadows with a simple color conversion rule.

The figures used for stimulation were created in the following rule. Only one component of RGB in the figure was changed to the maximum value (255). The drawings created by this method may be considered as colored shadow versions. We calculated the luminance (= M+L at DKL color space) of the brightest square and the darkest square and determined the contrast ratios. The lower contrast ratio was 0.04 at Figure S1(c) condition. However, the display color limit could not enable the contrast ratio of 0.04 for either red or blue. Thus, we selected the lowest contrast ratio as 0.13. As we separated the effect of color difference and contrast ratio, we selected three contrast conditions: 0.13, 0.27, 0.36. The hue was changed with a contrast and was rotated on the S- (M + L) vs. L-M plane in the DKL color space.

### Supplement 2 Magnitude estimation task and procedures in Experiment 1.

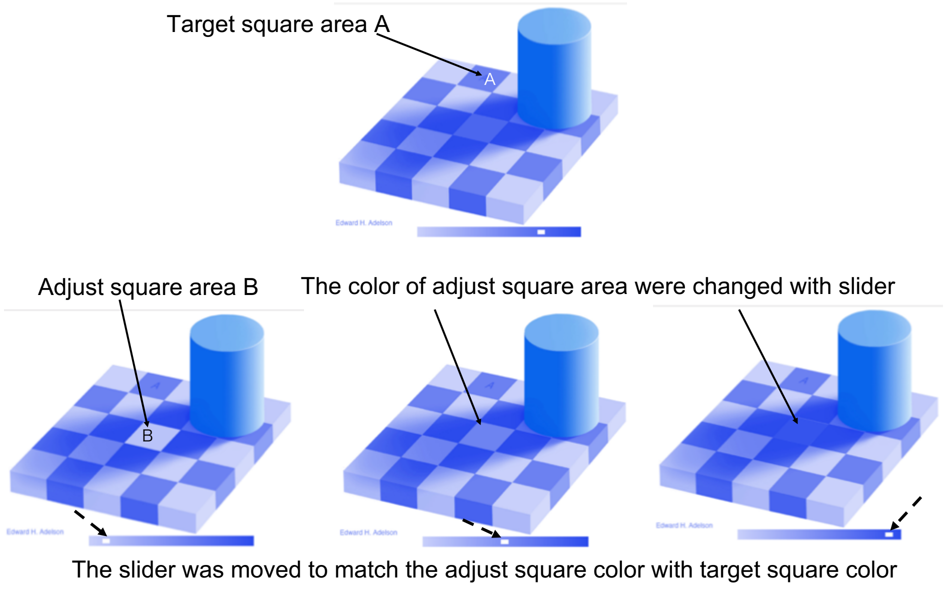


Figure S2. Typical example of our procedure in Experiment 1.

An example task in Experiment 1 is shown. Subjects move the slider (denoted by the dotted allow) on the bottom scale to change the color of square B in the cast shadow region. Subjects move the slider to adjust B to be the same brightness/color as square A, the target. When subjects perceive B and A to be the same brightness/color, they double click the mouse button. The amount that the slider is moved is used to indicate the illusion intensity. The stimuli were created in javascript and executed on the browser.

### Supplement 3 Shade stimuli of Experiment 2.

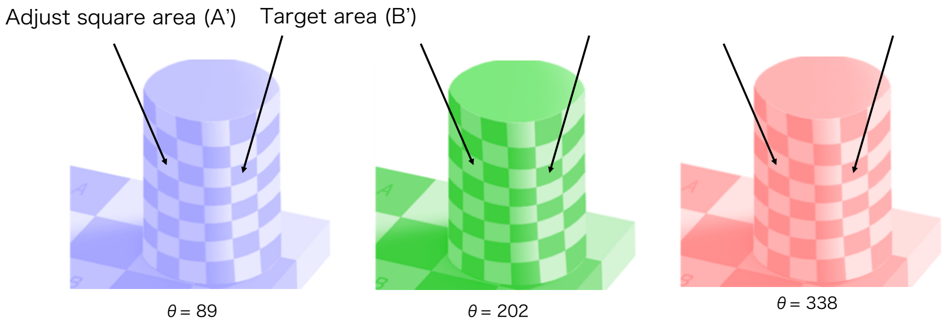


Figure S3. Examples of shade stimuli of Experiment 2.

We incorporated another illusion, the checker shade illusion, but with the same contrast and hue configurations. The shade illusion stimuli are shown in Figure S3. The numbers under each figure — 89, 202, and 336 — represent hues in the DKL color space. The cylinder was expanded because the area of the square was adjusted to be the same size as the cast shadow illusion.

**Supplement 4 Results of Experiment 1 with three contrast ratios.**

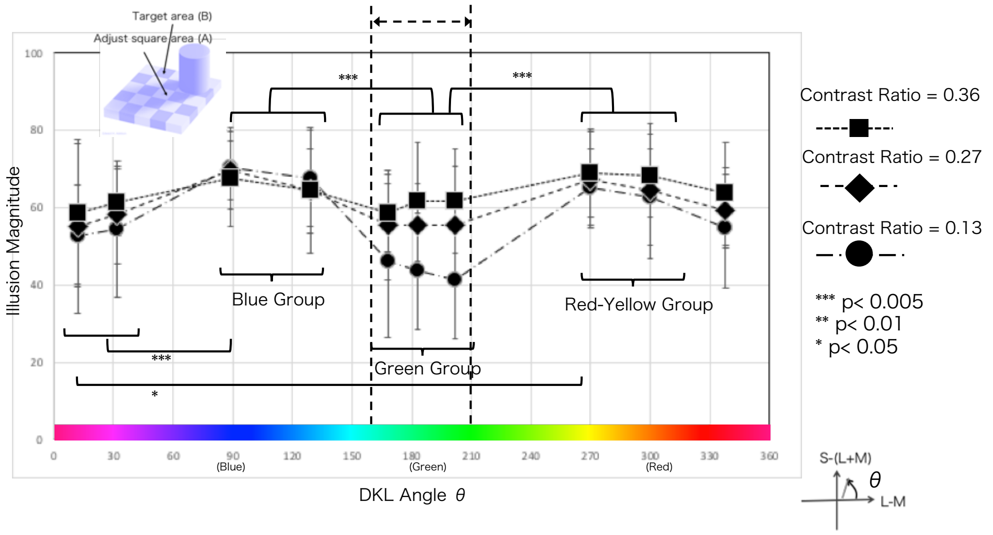


Figure S4. Results of experiment1; Cast Shadow illusions with different contrast levels. More detailed results are shown for the effects of contrast and hue. Two-way ANOVA indicated that the effects of both contrast and color were significant (main effect of color: $F(9, 17612) = 9.9730$, $P < 0.001$; main effect of contrast ratio: $F(2, 4389) = 11.1836$, $P < 0.001$) with interaction between these factors ($F(18, 4363) = 1.2353$, $P = 0.2281$). Pairwise comparisons using a Bonferroni Correction indicated significant differences in attenuation of illusion magnitude between the blue group and the green group and also between the red-yellow group and the green group. The color difference is related to attenuation in the illusion magnitude. A one-way ANOVA indicated that the effects of color with two contrast ratios were significant (main effect of color at contrast ratio = 0.13 : $F(9, 15197) = 7.9939$, $P < 0.0001$, color at contrast ratio = 0.25 : $F(9, 4427) = 2.479$, $P = 0.0109$), although there was a nonsignificant trend at contrast ratio = 0.36:$F(9, 2351) = 1.448$, $P = 0.173$).

At contrast conditions, pairwise comparisons using the Mann–Whitney U test indicated that there was a significant difference between contrast condition 0.13 and 0.36 ($P = 0.001$). There were no significant differences between either 0.13 and 0.27 ($P = 0.125$) or 0.27 and 0.36 ($P = 0.315$).

The attenuation of illusion magnitude obtained cannot be attributed to the luminance contrast difference. It should be noted that the effect is due to the difference in color. Because of the limitations of the rules making up Adelson's variation of the checker shadow, reducing the contrast ratio will increase the brightness. With increasing brightness, the color mode moves closer to the fluorescence mode, making it easier to estimate more colored ambient light.

### Supplement 5 Results of Experiment 2 with three contrast ratios.

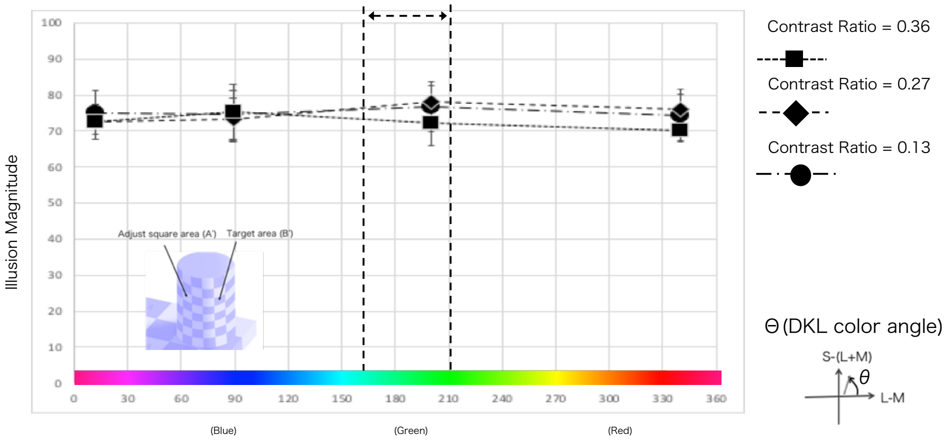


Figure S5. Results of Experiment 2: checker shade pattern with cylinder indicate which stimulus set and comparison were used; in this case, Figure S3.

More detailed results are shown for the effects of contrast and hue at the shade experience. Results of a two-way ANOVA for contrast ratio and color indicated significant main effects for both factors (color: $F(3,94) = 0.8281$, $P = 0.4812$; contrast ratio, $F(2,175) = 2.30684$, $P = 0.1045$) with an interaction between factors ($F(4,220) = 1.3662$, $P = 0.2530$). The findings revealed that neither color differences nor contrast ratios altered the illusion magnitude significantly. Moreover, the findings demonstrate that the contrast of shades is due to shape, and have no concerned with the color difference. The findings agreed well with the constant illusion magnitude.

**Supplement 6. The relationship between the direction of attenuation of illusion and natural scene color distribution in DKL color space.**

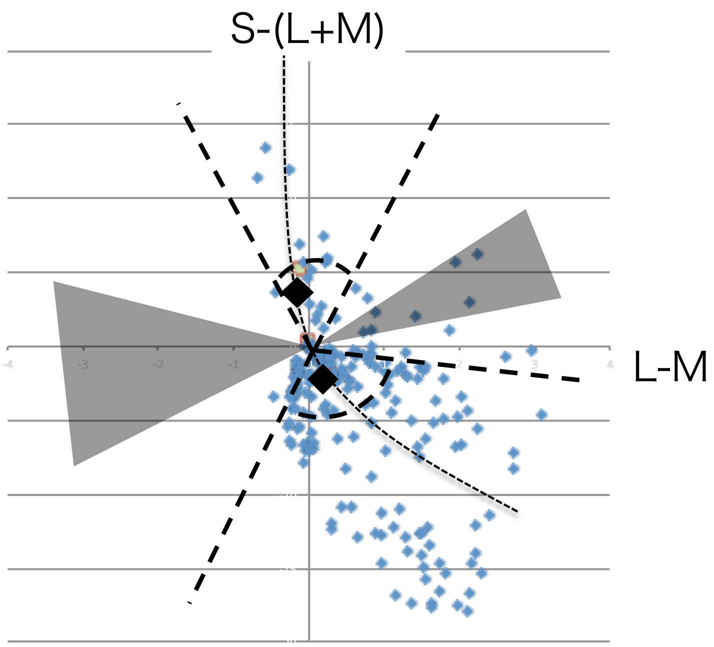


Figure S6-1 The attenuation light source color angles with sunlight trajectory and natural scene color statistics on L-M vs. S-(L+M) plane in DKL color space.

We hypothesized that, in an environment where illumination light is not presented explicitly, the object color is determined by the object color and the illumination color that maximizes the overlap with the color distribution of natural objects.

Figure S6-1 shows the attenuation color angles with sunlight trajectory and natural scene color statistics on L-M vs. S-(L+M) plane in DKL color space. The gray area shows the illusion attenuation angle. Two areas defined by broken lines indicate PCA of the spectrum change of sunlight in a day (judd). Two diamonds show two colors of #The Dress: brown and blue. From the statistical distribution of color and the positional relationship between diamonds, when the color green is considered as illumination, the overlap with the statistical distribution is increased by moving the diamond in the L direction (to the right in the figure). #The Dress shows that the actual perception assumes two colors, dark blue (shadow) and light yellow, which are consistent with the PCA so that the diamond will be shifted either up or down in Figure S6-1. The overlap between two diamonds and the natural color distribution was calculated using KL divergence as the overlap of probability distributions. KL divergence can measure the matching between two distributions. If the two distributions are the same, KL divergence = 0. The findings are presented using a solid line in Fig. 6-2. When two diamonds are maximum matched to the natural object statistical map, the direction in which the green illumination is taken into consideration has the smallest KL div at the dash-dot line in Figure S6-2. This result imposes a condition that the green region obtained in our experiments is difficult to estimate as natural illumination light at the dotted line in Figure S6-2. With the estimated probability of illumination light (dotted line in the figure) as a constraint, the KL divergence is minimized under conditions that take blue and light-yellow illumination light into consideration (solid line in Figure 6-2). Moreover, we assume that two-color points are stable because they are valley bottoms. In other words, in #The Dress type illusion, we can assume the same situation as when solving a transport problem as light source estimations become unstable. However, this transport problem is established only in a very narrow area, as the optimum value in multiple routes is limited. Two calculations of KL divergence are same Wassestein-Metric to solve the transport problem. In addition, the result of the sneaker that performed the same calculation is shown for reference in Figure S6-3.

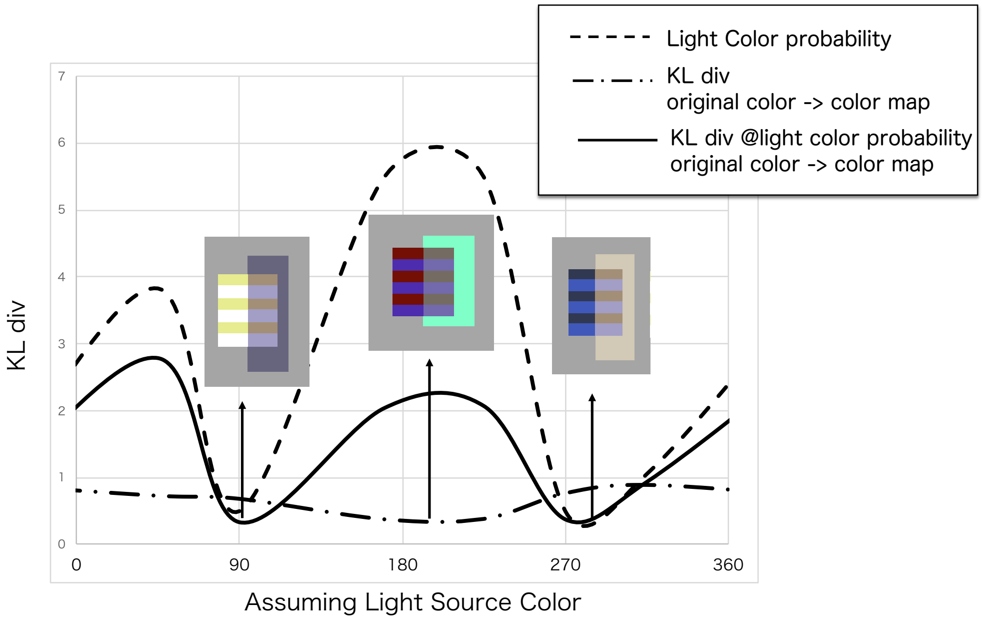


Figure S6-2 The results of calculation of KL divergence.
As #The Dress condition, two bottom are shown. In Figure, light and color combinations are shown at the mountain point and two bottom points. Note that KL divergence is small when the match rate is high. Combinations of two bottoms have a high probability of existence.

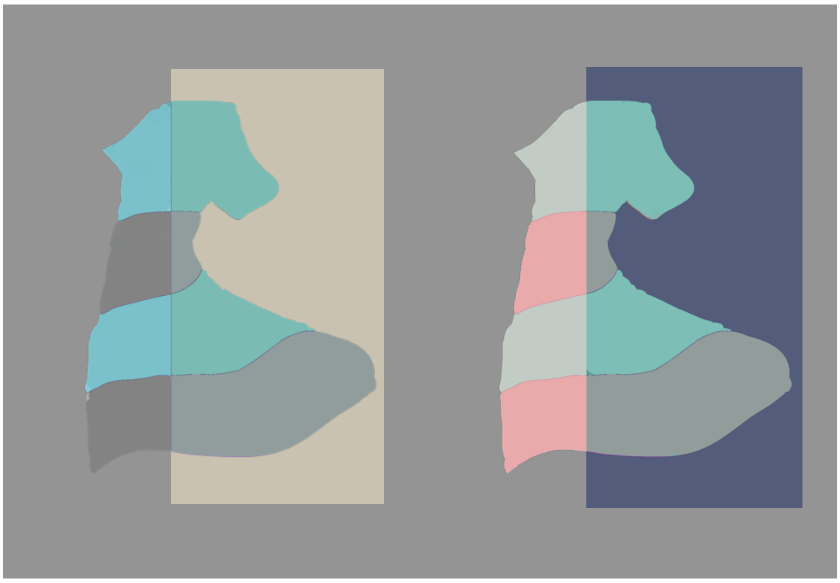

Figure S6-3. Color combination same as #The sneakers.

The color combination is also calculated with the conversion rule as same as the calculation rule of color conversion of #The dress.

### Supplement 7 Shade/Shadow illusion for several colors

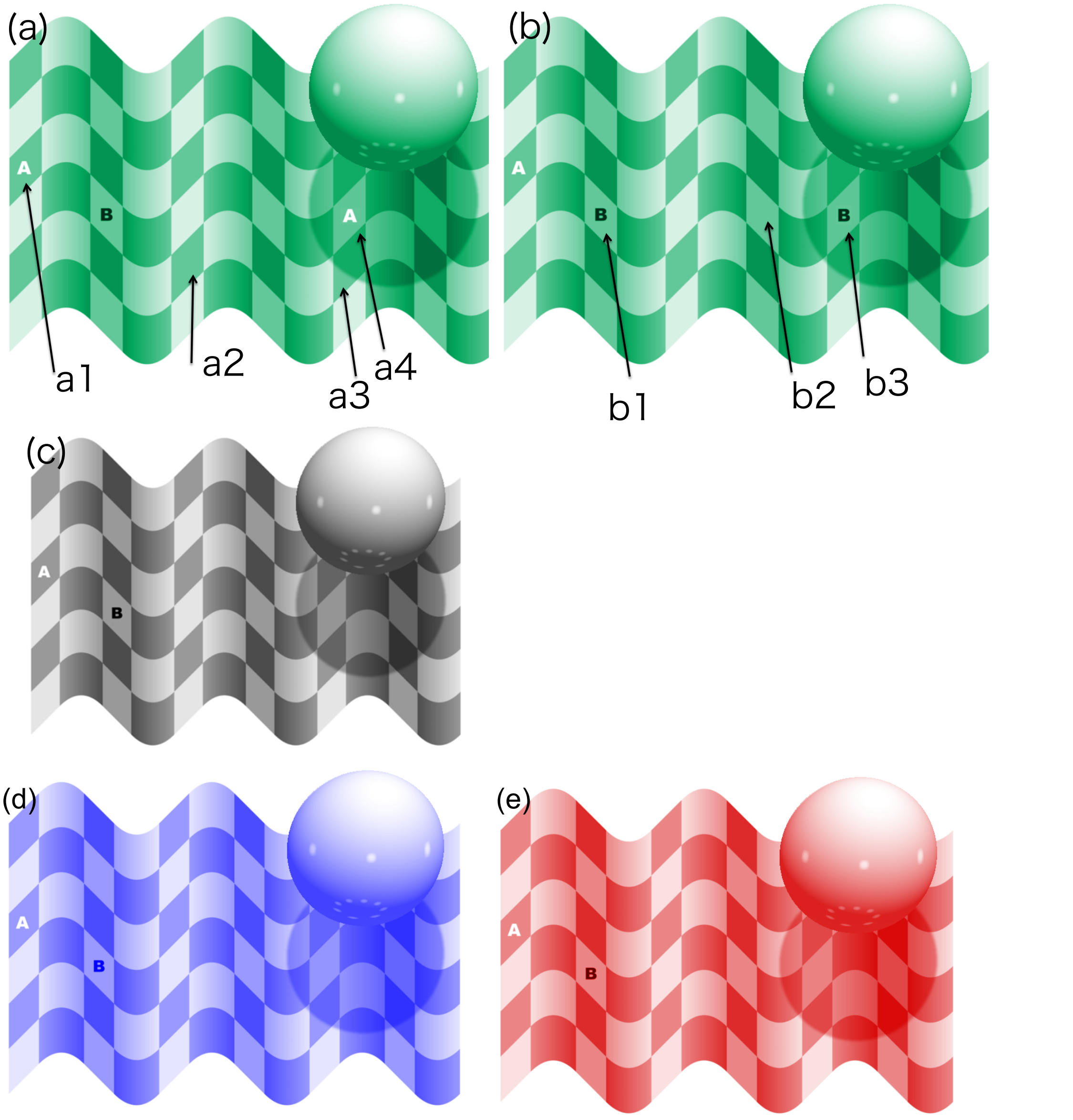


Figure S7. New illusion due to the different constancy characteristics; cast shadows and shades.

The difference in the brightness between squares A and B is essential. In Figure S7(a), a4 may seem brighter than a1. In that case, consider a3 and a4 to be the same brightness. In most cases, a3 and a4 can seem to be equally bright. a3 and a2 will appear to be equally bright. a2 and a1 are equally bright. Even in this case, a4 and a1 may not appear to be equally bright. However, it is easy to compare a4 and a3, a3 and a2, and a2 and a1. It is easy to compare b1, b2, and b3 in Figure S7(b). All will appear to be equally bright.

The square color a1 to a4 and b1 to b3 are all physically the same. The square color a4 and b3 are placed in the ball cast shadow. b1 and b2 are positioned in the shade. A white letter “A” in the square in Figure S7(a) is darker than the surroundings, and a black letter “B” is brighter than the surroundings, very weakly contrast bias is applied. Receiving the contrast bias, a4 appears to be equal in brightness to a3 without being affected by cast shadow. Meanwhile, b3 appears to be equal in brightness to b1 and b2. This, the difference between the cast shadow and the shade, is one of the new illusion examples created from the experimental results of this study. This phenomenon does not occur in Fig. S7(c), fabricated in monotone version. Figure S7(d), (e): Blue and Red color versions of the new type illusion. Only the green version is different. Only the green version can adjust a3 and a4 to the same brightness. In other color versions, a4 is always brighter than is a3. The target assigned is the same as in Figure S7(a).
